# Phage administration with repeated intravenous doses leads to faster phage clearance in mammalian hosts

**DOI:** 10.1101/2023.03.10.532150

**Authors:** Xin Tan, Kai Chen, Zhihuan Jiang, Ziqiang Liu, Siyun Wang, Yong Ying, Jieqiong Zhang, Shengjian Yuan, Zhipeng Huang, Ruyue Gao, Min Zhao, Aoting Weng, Yongqing Yang, Huilong Luo, Daizhou Zhang, Yingfei Ma

## Abstract

**Objectives:** Phage therapy has shown a great promise for the treatment of multidrug-resistant bacterial infections. However, the lack of a thorough and organized understanding of phage-body interactions has limited its clinical application.

**Methods:** Here, we administered different purified phages (*Salmonella* phage SE_SZW1, *Acinetobacter* phage AB_SZ6, and *Pseudomonas* phage PA_LZ7) intravenously to healthy animals (rats and monkeys) to evaluate the phage-induced host responses and phage pharmacokinetics (PK) with different intravenous (IV) doses in healthy animals. The plasma and the organs were sampled after different IV doses to determine the phage biodistribution, the phage-induced cytokines, and antibodies. The potential side effects of phages on animals were assessed.

**Results:** A non-compartment model revealed that the plasma phage titer gradually decreased over time following a single dose. Repeated doses caused that the plasma phage titer at 5 minutes dropped 2-3 Log_10_ compared to the first dose regardless of phage types in rats. Host innate immune responses were activated including the upregulated expression (>10-fold) of TNF-*α* and splenic enlargement following repeated doses. Phage-specific neutralization antibodies in animals receiving phages were detected. Similar results were obtained from monkeys.

**Conclusions:** The mammalian bodies were well-tolerant to the administered phages. The animal responses to the phages and the phage biodistribution profiles could have a significant impact on the efficacy of phage therapy.

## Introduction

The therapeutic potential of phages has been highlighted in numerous recent studies^1–6^. However, although no severe adverse effects of phage therapy in various animal models and clinical studies were observed ^4,7^, safety concerns have largely limited the use of phage therapy on a large scale. Phage particles consist of proteins and nucleic acids (DNA or RNA) that can induce innate and adaptive immune responses in the mammalian body^8,9^. Thus, phage-body interactions inevitably affect animal or human health, phage pharmacokinetics (PK), and the efficacy of phage therapy. However, research on phage-body interactions is limited. Clinical phage therapy studies frequently involve the concurrent use of other antimicrobial agents^1,2,10^. In addition, the majority of phage therapy studies were carried out in infectious animal models where pathogens and phage-mediated lysis of bacteria will release amounts of endotoxins and other proinflammatory products^4,11^. These investigations are unable to evaluate the full impacts of phages alone on the body. Furthermore, phages are among the most diverse biological entities on earth^12^. It remains unclear whether the effects of these diverse phages on the body are consistent. Therefore, it is urgently necessary to perform a comprehensive and systematic analysis of phage-body interactions with different phages under the same conditions.

Here, we address these issues by administering distinct highly purified phages to healthy animals by intravenous (IV) route as IV is the most effective and fastest way to deliver phages into the body, facilitating phage dissemination in all organs and tissues within minutes^8^. This study aims to evaluate the phage-induced host immune responses and PK of phages with different IV doses in healthy rats and monkeys.

## Methods

### Bacteria and Phages

*Salmonella typhimurium* strain SL7207 and *P. aeruginosa* strain PAO1 were from laboratory collections. *A. baumannii* clinical isolate was obtained from routine microbiological cultures of clinical samples. Three lytic phages including SE_SZW1 for *Samonella typhimurium* ^13^, AB_SZ6 for *Acinetobacter baumannii*, and PA_LZ7 for *Pseudomonas aeruginosa* (See details in **Supplementary Materials, Table S1** and **Figure S1**) were isolated from sewage water. See details in Supplementary Materials for phage purification. **Animals**

All animal experiments were approved by the Shandong Academy of Pharmaceutical Sciences Animal Ethics Committee. Male and female Sprague-Dawley rats (6-8 weeks old) and Male and female cynomolgus monkeys (5-7 years old) were used in this study. See **Supplementary Materials, Table S4, S5, S6, and S7** for more information, including the sex, weight, and age of the animals.

### PK Study

Healthy rats received phages SE_SZW1, PA_LZ7, and AB_SZ6 separately once daily by IV administration at 5*10^9^ (Low dosage, LD) or 5*10^10^ PFU/kg (High dosage, HD) for 7 d. Healthy monkeys received phage SE_SZW1 once daily by IV administration at 10^9^ PFU/kg (Relative low dosage, RLD) or LD for 14 d. Blood were collected at 5 and 30 min, and 1, 2, 4, 6, and 24 h following the first administration (**Figure 1A and Figure 5A**); at 5 min following 3, 5, and 7 IV doses in rats (**Figure 2A**); at 5 min following 3, 7, 11, and 14 IV doses in monkeys for phage enumeration (**Figure 5A**). Regression analysis on the phage titer in the plasma over time was performed using non-compartmental analysis. Rats were killed at 1, 24, or 72 h after a single dose with each phage; rats were killed at 1, 24, or 72h after 3 doses with phage SE_SZW1, and tissues were harvested and homogenized in SM buffer for phage enumeration. Phage enumeration was performed with the plaque assay and the qPCR method^14^.

**Figure 1.**
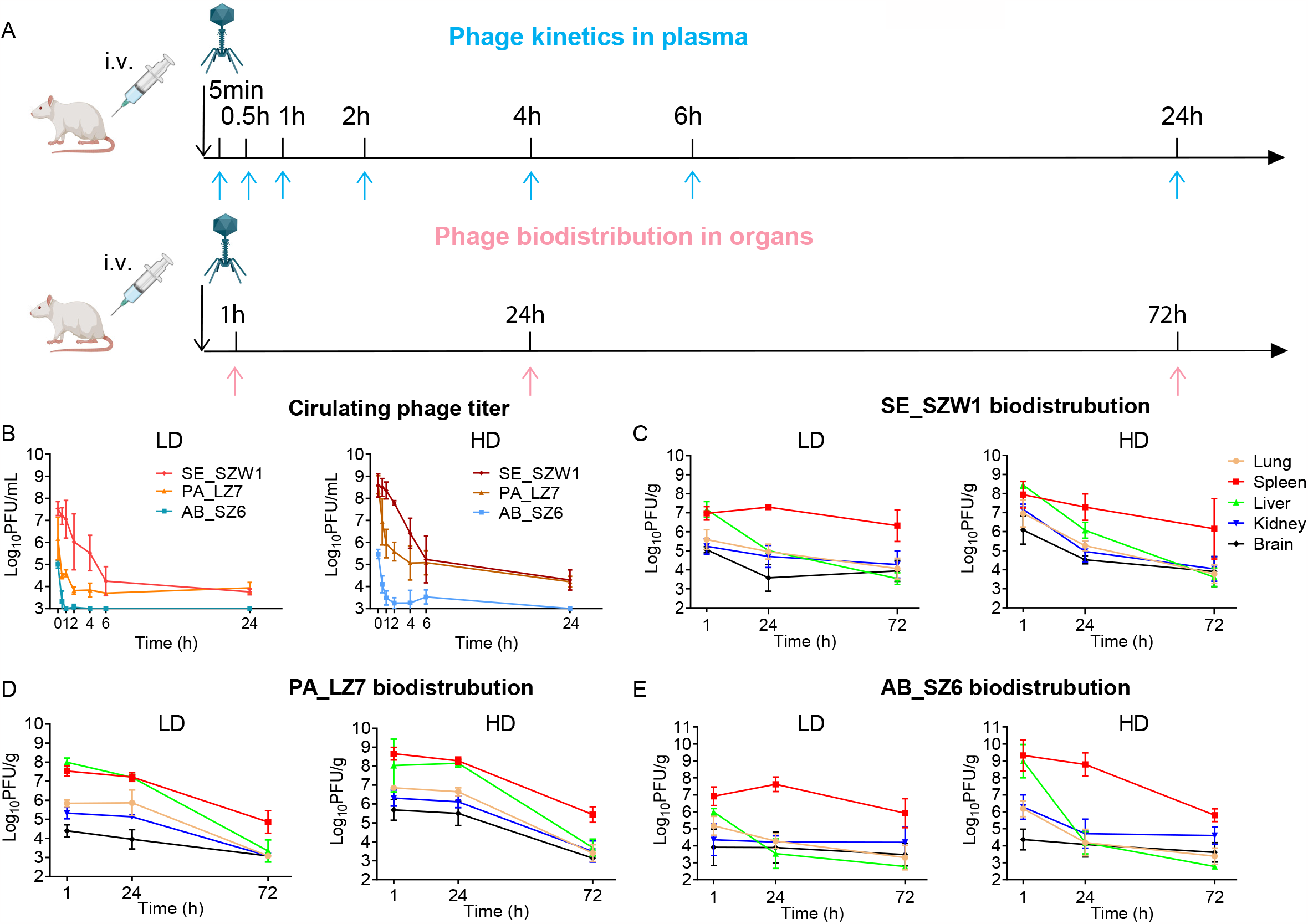
PK and biodistribution of phages with a single dose in rats. (A) Schematic representation of the experimental design. Phage kinetics in the plasma and phage biodistribution in organs were performed following a single dose administration in rats. (B) Kinetics of active phages SE_SZW1, PA_LZ7 and AB_SZ6 in the plasma following single intravenous (IV) administration at a dosage of 5*10^9^ (LD) and 5*10^10^ PFU /kg (HD) in healthy rats. Phage titer was obtained by plaque assay, phage titer is expressed as PFU per mL in the plasma, the lower limit of quantification (LLOQ) for phage SE_SZW1 and PA_LZ7 is 5000 PFU/mL and for AB_SZ6 is 1000 PFU/mL. Biodistribution of active phages SE_SZW1 (C), PA_LZ7 (D) and AB_SZ6 (E) in organs following single IV administration in healthy rats. Active phage titer is expressed as PFU per g of each organ, the LLOQ is 1200 PFU/g; phage titer was determined by plaque assay, and each symbol represents the means with standard deviation (sd).

**Figure 2.**
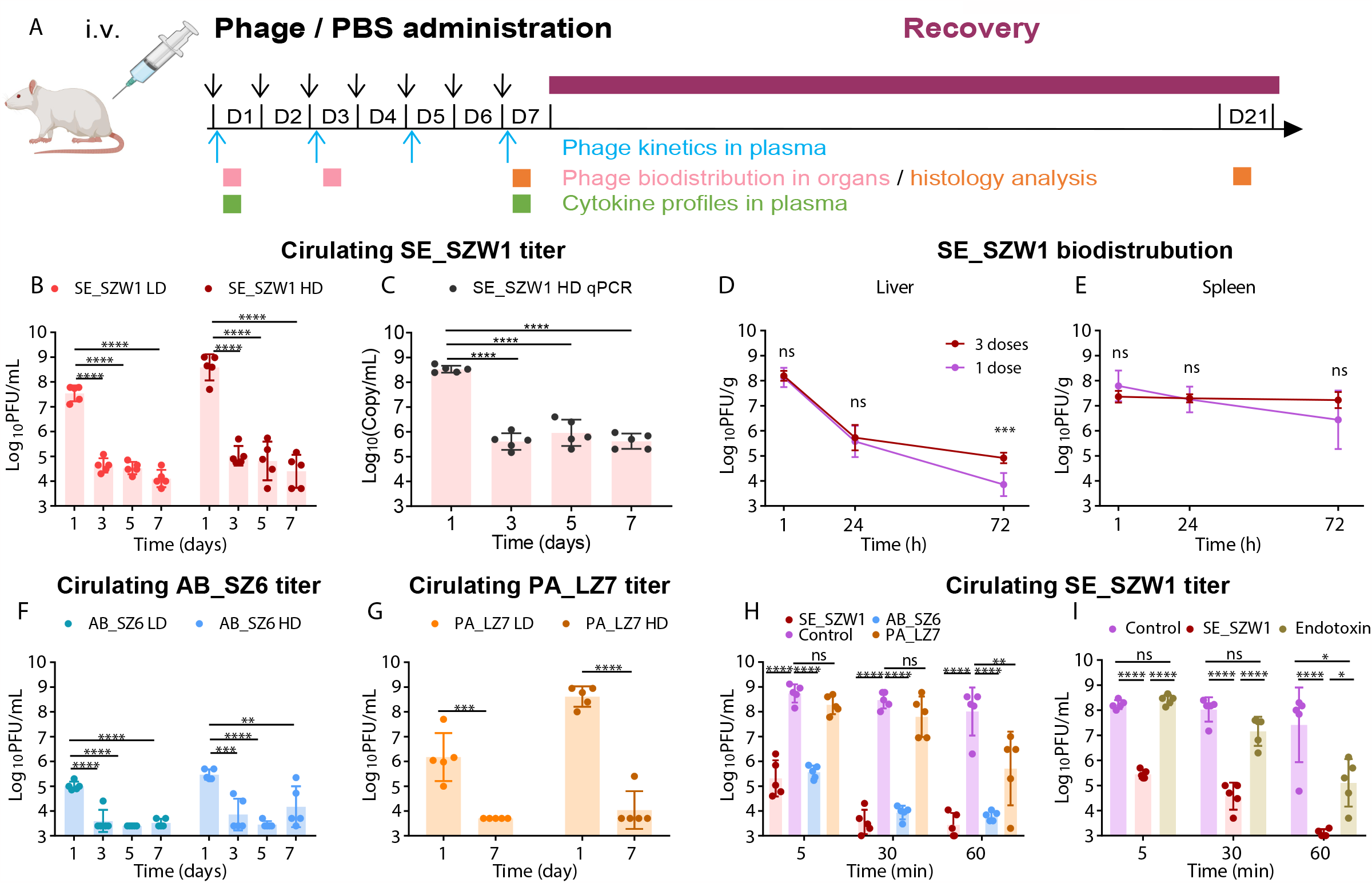
PK and biodistribution of phages with multiple doses in rats. (A) Schematic representation of the experimental design. Phage kinetics in the plasma, phage biodistribution in organs and histology analysis were performed following different doses of administration in rats. (B) Phage titer at 5 minutes in the plasma after different IV injections with phages SE_SZW1 in rats were presented. Phage titer was obtained by plaque assay, each symbol represents the means with sd (n=5); the LLOQ is 5000 PFU/mL. (C) Phage genome titer at 5 min in the plasma after different IV injections with phage SE_SZW1 in rats, phage genome titer is expressed as copy per mL; phage titer was determined by the qPCR method, each symbol representing the means with sd (n=5). Biodistribution of active phage in the liver (D) and spleen (E) after administration with 3 doses of phages SE_SZW1 HD (n=5) or a single dose (n=10). Active phage titer is expressed as PFU per g of each organ, the LLOQ is 1000 PFU/g. Phage titer was determined by plaque assay, each symbol representing the means with sd (ns: no significance; *** *P*<0.001). Phage titer at 5 minutes in the plasma after different IV injections with phages AB_SZ6 (F) and PA_LZ7 (G) in rats were presented. Phage titer was obtained by plaque assay, each symbol represents the means with sd (n=5); the LLOQ is 5000 PFU/mL. (H) Phage SE_SZW1 kinetics in plasma after the administration with phage SE_SZW1 HD in rats pretreated with 2 doses of phages SE_SZW1 HD, AB_SZ6 HD, PA_LZ7 HD or PBS. Phage titer was obtained by the plaque assay, and each symbol represents the means with sd (n=5). The LLOQ is 1000 PFU/mL (**, *P*<0.01; *** *P*<0.001; **** *P*<0.0001; ns: no significance). (I) Phage SE_SZW1 kinetics in the plasma after administration with phage SE_SZW1 HD from animals pretreated with 2 doses of phages SE_SZW1 HD, *Salmonella* endotoxin or PBS. Endotoxin was diluted with PBS to the same level of SE_SZW1 HD group. Phage titer was obtained by plaque assay, each symbol represents the means with sd (n=5). The LLOQ is 500 PFU/mL (**, *P*<0.01; *** *P*<0.001; **** *P*<0.0001).

### Cytokine Analysis

Blood were collected at 1, 6, and 24 h following phage administrations on Day (D) 1 and D7 in rats (**Figure 2A**); on D1 and D14 in monkeys for cytokine analysis (**Figure 5A**). The cytokines level of rats was evaluated by the V-PLEX Proinflammatory Panel 2 Rat Kit and the cytokines level of monkeys was measured by the Elisa Kit following the manufacturer’s instructions.

### Adaptive Immune Response Study

Following phage administration, animals underwent a 14-day recovery period. Plasma were collected before on D1, 3, 5, 7, 10, 12, 15, and 21 with phages SE_SZ1 and AB_SZ6 in rats (**Figure 4A**); on D1, 3, 7, 11, 14, 21 and 28 with phages SE_SZ1 in monkeys (**Figure 5A**) for the phage neutralization assay and the Western blot analysis using the previously described method with several modifications^15^. See details in **Supplementary Materials**.

### Tolerance Study

A tolerance study was performed using the previously described method with several modifications^16^. See details in **Supplementary Materials**.

### Data analysis

Comparisons were performed by one-way ANOVA with the Bonferroni’s multiple comparison test and Student’s t-test except for special explanation. All statistical analyses were performed using Prism 7.04 (GraphPad, San Diego, CA, USA), and differences with *P*< 0.05 were considered statistically significant.

## Results

### PK and Biodistribution of Phages with A Single Dose in Rats

Following a single IV dose of phages, **Figure 1B** showed the concentration-time profiles of active phages. Overall, the plasma active phage titer gradually decreased over time regardless of the administered dosage. Nevertheless, these three different phages demonstrated distinct PK patterns in rats. The non-compartmental analysis indicated that the AUC_inf_ exposure was 5.27±0.13 Log_10_(h*PFU/mL) for phage AB_SZ6, 8.23 ± 0.24 Log_10_(h*PFU/mL) for phage PA_LZ7 and 8.81±0.22 Log_10_(h*PFU/mL) for phage SE_SZW1 in the HD groups. More PK parameters are shown in **Table S2**. Phage SE_SZW1 showed the slowest clearance, followed by phage PA_LZ7 and phage AB_SZ6 in the HD group. At 5 min after IV administration, the titer of phage SE_SZW1 was 7.54 ± 0.32 and 8.59 ± 0.53 Log_10_PFU/mL in the LD group and HD group, respectively. At 24 h, the titer of phage SE_SZW1 dropped by about 4 log_10_PFU/mL to 3.76 ± 0.13 and 4.29 ± 0.46 Log_10_PFU/mL in these two groups (**Figure 1B**). Interestingly, the healthy rats showed drastically fast clearance of phage AB_SZ6 (**Table S2**) from the blood. At 5 min after IV administration, the titer of phage AB_SZ6 decreased to about 2 Log_10_PFU/mL, lower than those of other two phages (**Figure 1B)**. Similar findings were observed in the LD group (**Table S2**).

These three phages demonstrated a similar distribution pattern that all phages primarily accumulated in the spleen and liver at 1 h post-administration and gradually decreased in all organs **(Figure 1C, 1D, and 1E)**. Then, the active phage titer in all organs decreased globally within 72 h, albeit it remained noticeably higher in the spleen than in other organs. For example, following a single dose IV phage (SE_SZW1) administration, at 1 h post-administration, the spleen and liver had substantially higher active phage titer (6.97 ± 0.35 Log_10_PFU/g, 7.14 ± 0.42 Log_10_PFU/g, *P*<0.0001) than the lung (5.59 ± 0.52 Log_10_PFU/g), kidney (5.24 ± 0.41 Log_10_PFU/g) and brain (5.06 ± 0.16 log_10_PFU/g) in LD group **(Figure 1C)**; at 72h, the active phage titer in the spleen remained at 6.32 ± 0.84 Log_10_PFU/g, whereas the active phage titer in the liver and other organs dropped to 3∼4 Log_10_PFU/g **(Figure 1C)**. In addition, the biodistribution patterns of phage SE_SZW1 in the HD group obtained from the quantitative PCR analysis showed similar results (See more details in **Supplementary Materials** and **Figure S2**).

### PK and Biodistribution of Phages with Repeated Doses in Rats

To investigate the dynamics of the active phage titer in rats with multiple IV doses, we measured the phage concentration in plasma following 3, 5, and 7 repeated IV phage doses (**Figure 2A**). We observed an overall decrease of active phage titer following repeated doses for all these phages compared to the first dose. For example, the active phage titer of phage SE_SZW1 decreased sharply (*P*<0.0001) in plasma by 3 magnitudes at 5 min after 3 IV doses compared to that of the first dose in both LD and HD groups (**Figure 2B**) and the active phage titer in plasma after 5 and 7 repeated doses remained at an extremely lower level (*P*<0.0001) than that of the first dose as well. This observation was supported by the qPCR assay in the HD groups as well (**Figure 2C**). However, we did not observe enhanced phage clearance within 72 h in the spleen and liver following 3 doses compared to a single dose (**Figure 2D and 2E**). The titer of the active phages AB_SZ6 and PA_LZ7 at 5 minutes post-administration in plasma dropped drastically (*P*<0.01) following repeated IV dosing as well in a manner similar to that of phage SE_SZW1 (**Figure 2F and 2G**). We further observed that this enhanced phage clearance in rats caused by repeated phage administration was non-specific. The phage SE_SZW1 kinetics were measured after the phage SE_SZW1 administration in rats pretreated with 2 doses of phages SE_SZW1, PA_LZ7, AB_SZ6 or phosphate-buffered saline (PBS, Control). At 5 min after the phage SE_SZW1 administration, the active phage titer in plasma was ∼3 Log_10_PFU/mL lower (*P*<0.0001) in the rats pretreated with 2 previous phage doses of AB_SZ6 compared to the rats of control; at 1 h, the active phage titer was significantly decreased (*P*<0.01) in the rats pretreated with phage PA_LZ7 compared to the rats of control (**Figure 2H**). A significant enhanced (*P*<0.05) phage clearance was also observed in the rats pretreated with *Salmonella* endotoxin (187.5 EU/kg, same to that in the phage SE_SZW1 preparation (HD)) (**Figure 2I**).

### Cytokine Analysis in Rats

We investigated the host innate immune responses following phage IV administration. We measured cytokine concentrations in the plasma at 1 and 24 h following phage administrations on Day (D) 1 and D7 for these three phages (**Figure 2A**). We observed obvious alterations in TNF-*α*, IL-6, and KC/GRO (**Figure 3 and S4**). These cytokines altered in a dosage-dependent manner, *i*.*e*. the HD group rats had a higher level of cytokines than the LD group rats, and returned to a normal range within 24 h. The alteration of these cytokines became milder following 7 doses than the first dose. For example, in the case of phage SE_SZW1, at 1 h post-administration of the first dose, the concentration of TNF-*α* increased by 80-fold (*P*>0.05) in the LD group, but remarkably increased (*P*<0.001) by 300-fold in the HD group compared to that of the control group (**Figure 3A**). The concentrations of IL6 and KC/GRO responded similarly but the effect was mild. Following 7 doses, the concentration of TNF-*α* significantly increased in the LD (*P*<0.0001) and HD groups (*P*<0.001), but was much milder than that of the first dose group (**Figure 3A**). This pro-inflammatory response following phage administration was similar to that of residual endotoxin administration (**Figure S4**). The changes in cytokines profiles following IV administration for phages AB_SZ6 and PA_LZ7 were similar to that of phage SE_SZW1 **(Figure 3B and 3C**). These results confirmed that IV phage administration can induce host innate immune response in rats.

**Figure 3.**
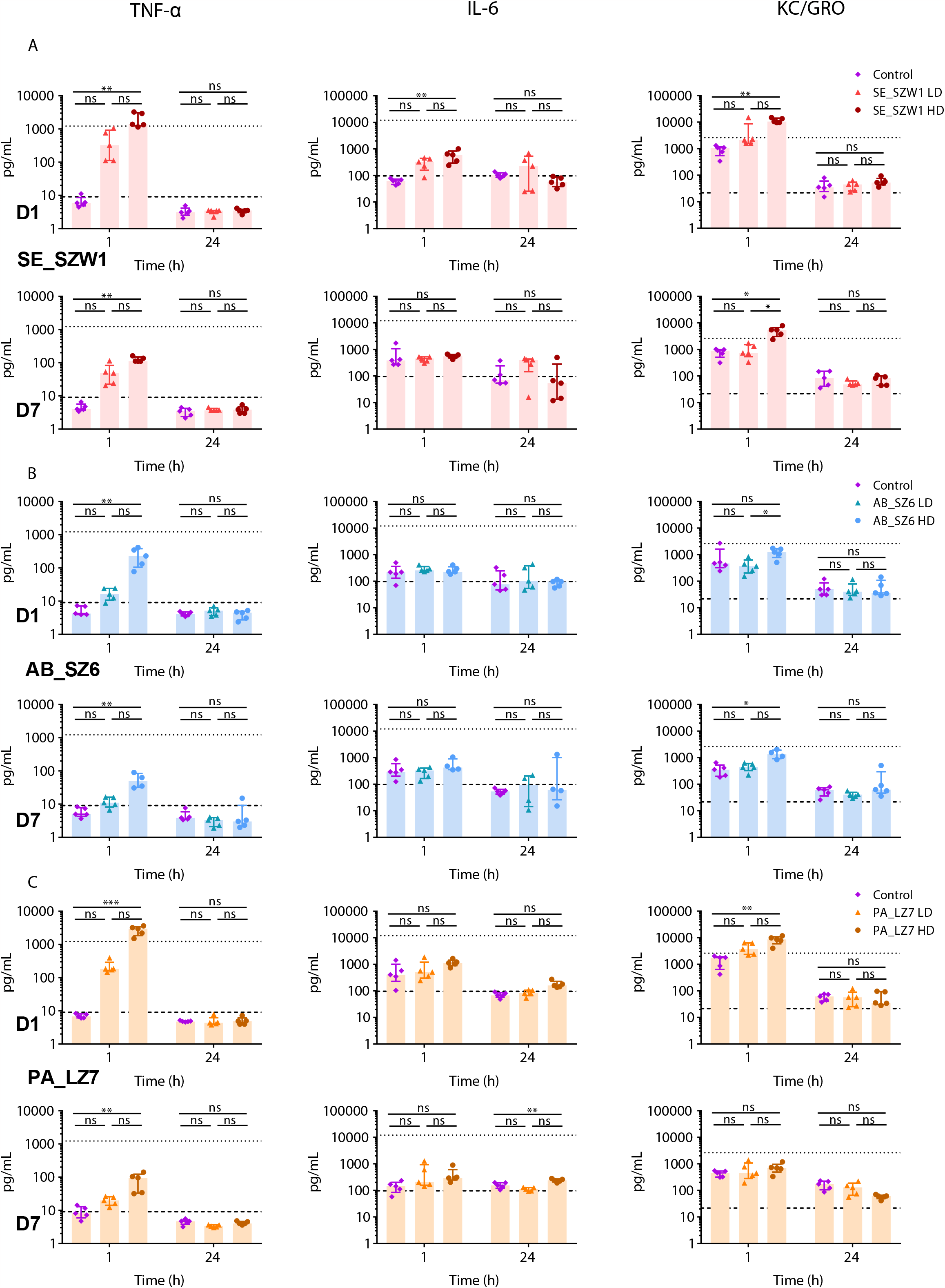
Cytokine profile in the plasma of rats. Cytokine concentrations in the plasma for rats were measured at 1 and 24 h with phage SE_SZW1 (A), AB_SZ6 (B) and PA_LZ7 (C) following a single dose or 7 repeated doses on D1 and D7. Data are presented as median with the interquartile range (IQR). The dashed line represents as the LLOQ and dotted line represents the upper limit of quantification (ULOQ) on the graph. For those points below LLOQ and above ULOQ, assign the value of 1/2 LLOQ and 2 ULOQ, respectively, for statistical analysis. Comparisons performed exclusively within the same time point (n=5; *, *P*<0.05; **, *P*<0.01; *** *P*<0.001; ns: no significance). Analysis was performed by Kruskal-Wallis with Dunn’s test.

### Adaptive Immune Responses Study in Rats

We also observed phage-induced adaptive immune responses in rats. The plasma had substantial anti-phage activity (*P*<0.0001) after 15 days post-administration of phage SE_SZW1 (**Figure 4B**). The plasma after 21 days post-administration of phage SE_SZW1 had the strongest anti-phage neutralization activity (*P*<0.0001) and reduced the active phages by about 3 Log_10_PFU/mL in the HD group rats (5 out 5 rats) (**Figure 4B**) while the plasma in the LD group rats had a mild phage-neutralization activity. This case was similar to that of phage AB_SZ6 (**Figure 4C**). This neutralization activity was phage-specific (**Figure 4D**). Western blot analysis using the sample of one rat from the HD group with phage SE_SZW1 indicated that strong IgG antibody recognition to the tail tube protein (25 kDa) and tail protein (93 kDa) with increasing signal over the experiment course (**Figure 4E**). Strong bindings to tail tube proteins were also observed in other rats on D21 (**Figure 4E**). Furthermore, WB analysis showed strong IgG antibody recognition to the capsid subunits (37 kDa) of phage AB_SZ6 on D21 (3 out of 5 rats) (**Figure 4F)**. This indicated that different phages had different immunogenicity.

**Figure 4.**
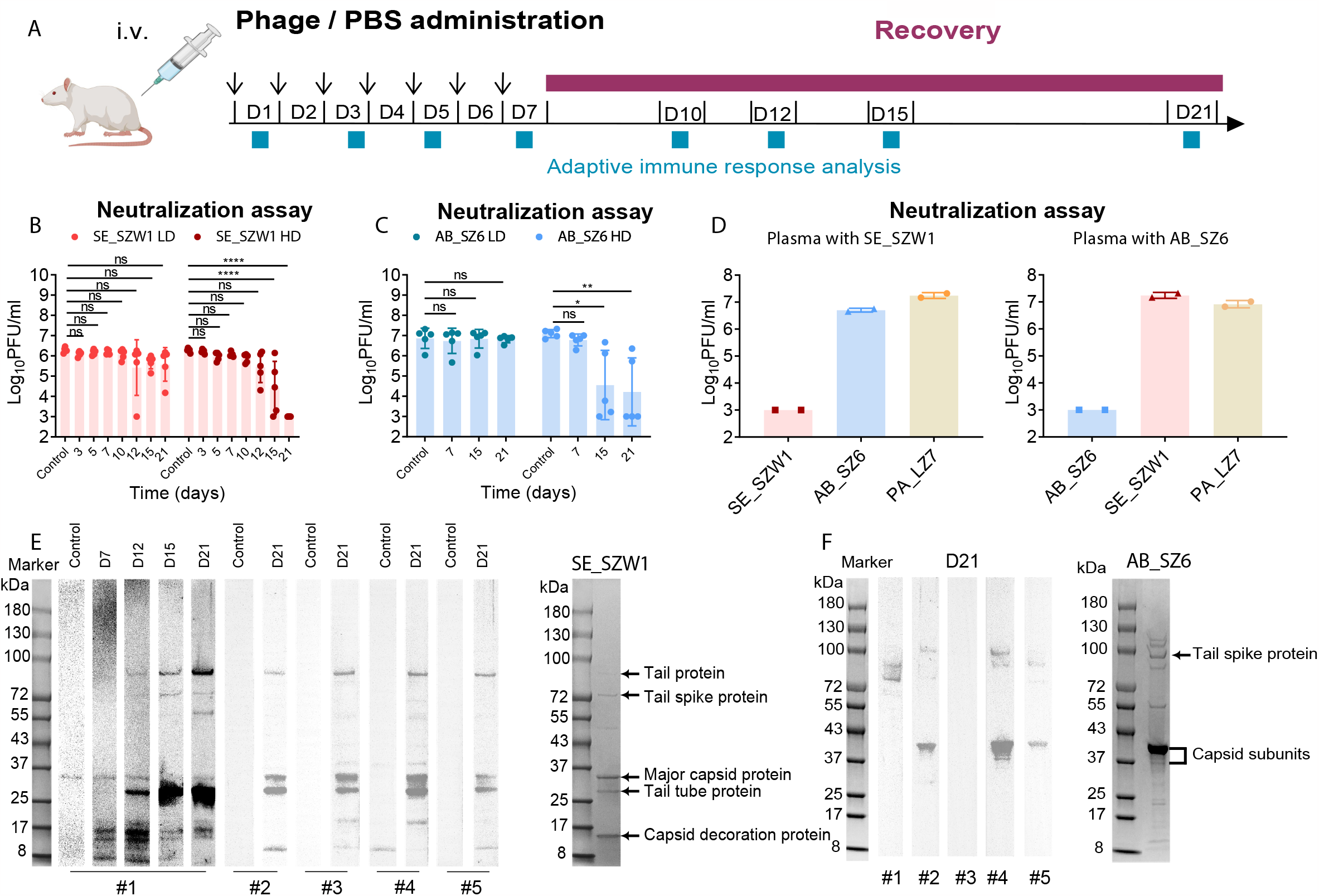
Adaptive immune responses study in rats. (A) Schematic representation of the experimental design. Plasma on D1, 3, 5, 7, 10, 12, 15, and 21 were collected for phage neutralization assay and western blot analysis for animals that received phage SE_SZ1 and AB_SZ6. Phage neutralization assays, where the plasma from rats with IV injections of phage SE_SZW1 (B) and AB_SZ6 (C) was incubated with phage (4.5*10^5^ PFU phage) for 24 h, then serially diluted 10-fold and plated on lawns of its host. The plasma of rats before phage treatment (D1) was set as control. The plasma of rats post-treatment until D21 is shown. Data are presented as means with sd. Comparisons performed for time points post-treatment to the control (n=5; *, *P*<0.05; **, *P*<0.01; *** *P*<0.001; **** *P*<0.0001; ns: no significant). (D) Phage neutralization assays, where plasma from rats of HD groups of phage SE_SZW1 or AB_SZ6 were incubated with phage SE_SZW1, AB_SZ6 or PA_LZ7 for 24 h, then serially diluted ten folds and plated on lawns of their corresponding host. Plasma with phage SE_SZW1 or AB_SZ6 post-treatment D21 is chosen (n=2 for each group). The LLOQ is 1000 PFU per mL. Western blot analysis of plasma responses to the phage SE_SZW1 (E) and AB_SZ6 (F) using 1:1000 plasma dilutions (as indicated) and detection with IgG-specific secondary antibodies in rats. The number # refers to the individual rat.

### PK of Phage SE_SZW1 and Phage-Induced Immune Responses in Monkeys

We conducted an experiment in nonhuman primates (NHPs) using phage SE_SZW1 to investigate whether the results are consistent across animal species and whether these findings can be translated into humans. To reduce the potential effect of endotoxin, we adjusted the phage dosage to 10^9^ (Relative low dose, RLD) and 5*10^9^ PFU/kg with endotoxin levels (with no detrimental effect for humans) at 3.75 and 18.75 EU/kg, respectively. While a minor difference in the PK profile of monkeys with a single dose was observed compared to that of rats (**Figure 5B and Table S3**), both groups exhibited a significant (*P*<0.05) decrease in active phage titer at 5 min after 3 repeated IV doses compared to the first dose (**Figure 5C**). The active phage titer remained at the LLOQ level after 7, 11, and 14 repeated doses.

**Figure 5.**
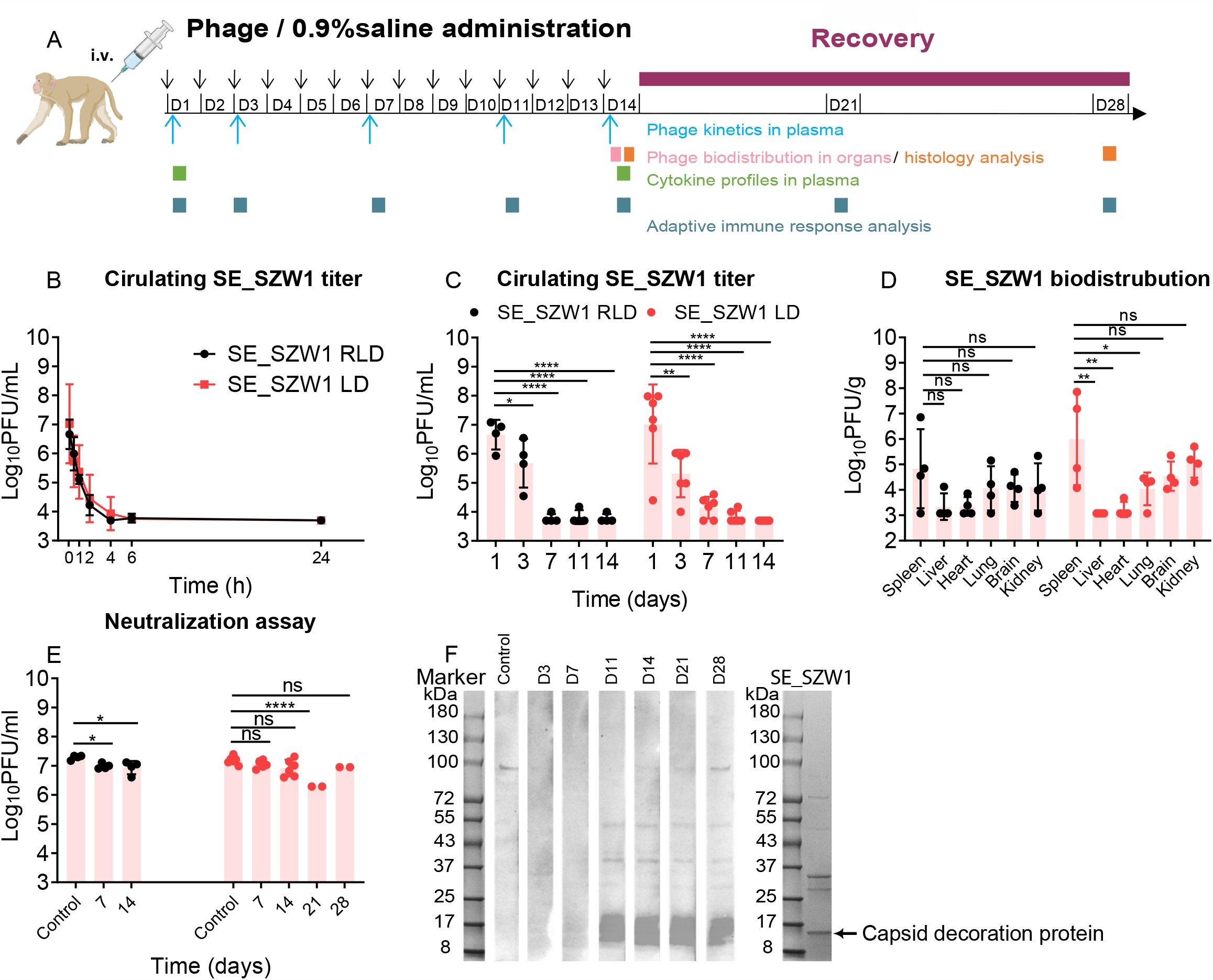
PK of phage SE_SZW1 and phage-induced immune responses in monkeys. (A) Schematic representation of the experimental design. Phage kinetics in the plasma, phage biodistribution in organs, cytokine response, histology analysis, and phage-specific antibody response in the plasma were performed following different dose administrations in monkeys. (B) Kinetics of phage SE_SZW1 in plasma following first IV administration at dose 10^9^ (RLD) and 5*10^9^ PFU/kg (LD) in cynomolgus monkeys. (C) Active Phage titer at 5 minutes in the plasma after different IV injections with phage SE_SZW1 in monkeys. Phage titer is expressed as PFU per mL in the plasma, phage titer was obtained by plaque assay, each symbol represents the means with sd (RLD group: n=4; LD group n=6), and the LLOQ is 5000 PFU/mL. (D) Biodistribution of phage SE_SZW1 in organs following 14 IV injections in monkeys. Phage titer is expressed as PFU per g of each organ, phage titer was determined by plaque assay, each symbol represents the means with sd (n=4), and the LLOQ is 1200 PFU/g (*, *P*<0.05; **, *P*<0.01; *** *P*<0.001; **** *P*<0.0001; ns: no significant). (E) Phage neutralization assays, where plasma from monkeys with IV injections of phage SE_SZW1 was incubated with phage (4.5*10^5^ PFU) for 24 h, then serially diluted 10-fold and plated on lawns of its host. The plasma of rats before phage treatment (D1) was set as control. The plasma of monkeys post-treatment until D28 is shown. The LLOQ is 1000 PFU per mL. Data are presented as means with sd. Comparisons performed for time points post-treatment to the control (RLD group: n=4; LD group n=6 except at D21 and 28 n=2). (*, *P*<0.05; **, *P*<0.01; *** *P*<0.001; **** *P*<0.0001; ns: no significant). (F) Western blot analysis of plasma responses to the phage SE_SZW1 using 1: 1000 plasma dilutions and detection with IgG-specific secondary antibodies in one monkey.

Animals were sacrificed at 24 h following 14 doses (**Figure 5A**), and we observed that phages primarily accumulated in the spleen, with phage titers of 6.00 ± 1.81 log_10_PFU/g and 4.83 ± 1.55 Log_10_PFU/g for RLD and LD groups, respectively (**Figure 5D**). Cytokines profiles showed no significant changes during the experiment for both groups (**Figure S5**). The neutralization assay revealed a significant (*P*<0.05) anti-phage effect in the plasma on D7 and D14 for RLD group and on D21 for LD group, respectively (**Figure 5E**). Moreover, a robust IgG antibody recognition to the capsid decoration protein was observed on D11 while the signals to major capsid, tail tube protein, and tail protein were very weak (**Figure 5F)**. This result was different from the profile of rats.

### Tolerance Study of Phages with Repeated Doses in Healthy Animals

Increased (*P*<0.0001) relative weight of the spleen was observed after 7 doses for all three phages in the HD group in rats, but not observed following 14-day recovery (**Figure S6**). Slightly extramedullary hematopoiesis was observed in the spleen samples after 7 doses (**Figure S7**) and recovered after 14-day recovery. No other toxicity effects were observed for these animals (See details in **Supplementary Materials**).

## Discussions

In this study, we evaluated the PK and phage-induced host immune responses of three different phages in healthy animals. We revealed a substantial and significant drop of phage titer in the plasma following repeated IV doses in both rats and monkeys regardless of phage types or dosages (**Figures 2, 5**, and **S3**). Our findings demonstrated that the administration of these three phages induced both innate immune responses and adaptive immune responses that produced phage-specific neutralizing antibodies. These immune responses and biodistribution profiles could have a significant impact on the efficacy of phage therapy.

Our results revealed the enhanced non-specific phage clearance following repeated phage administrations in animals (**Figures 2, 5 and S3**). However, the conclusion was limited by the fact that we can not completely eliminate bacterial endotoxin from the phage preparation. Further, we found that *Salmonell* endotoxin alone can also induce non-specific clearance (*P*<0.05) and the level was lower (*P*<0.05) than that by the phage SE_SZW1 preparation (**Figure 2I**). The enhanced phage clearance following repeated IV phage doses at 2*10^7^ PFU/kg (residual endotoxin level: ∼0.08 EU/kg)(**Figure S3**) also was observed. Thus, these results suggested that both endotoxin and phages played important roles in the enhanced phage clearance.

Phage clearance has been reported to be mostly attributable to the phagocytosis of phage particles in the spleen and liver^8,17–19^. However, we found no enhanced phage clearance in the spleen and liver following repeated IV doses (**Figure 2D and 2E**). This result suggested that more factors likely were involved in the enhanced phage clearance in plasma after repeated doses. The clear mechanism involved in this process needs further investigation.

In conclusion, our study revealed faster phage clearance with repeated IV doses in rats and monkeys and demonstrated the potential effect of host immune responses on phage clearance. Therefore, it is urgently necessary to conduct investigations for various phage formulations, including encapsulated phage to address the issues of IV phage administration raised by this study.

## Acknowledgments

We thank Dr. Zhang Wang from South China Normal University for his critical review of the manuscript. We thank Shengkun Dai from Shenzhen Institutes of Advanced Technology for performing mass spectrometry. This work received support from the Strategic Priority Research Program of the Chinese Academy of Sciences (No. XDB29050501); Guangdong Provincial Key Laboratory of Synthetic Genomics (No. 2019B030301006); Shenzhen Institute of Synthetic Biology Scientific Research Program (No. JCHZ20200001); National Natural Science Foundation of China (No. 32001038); Shenzhen Outstanding Scientific Innovation Talents Training Project (No. RCBS20210706092214015); Programs Foundation of 20 articles for colleges and universities in Jinan (No. 2020GXRC026).

## Author contributions

X.T., K.C. and Y.M. conceived and designed the experiments. X.T., K.C., and Y.M. supervised the project. Z.J., Y.Y., S.W., Z.L, M.Z, J.Z., Z.H., R.G., S.Y. and A.W. performed the experiments. All authors analyzed and discussed the data. X.T. wrote the original draft, with Y.M. providing further feedback and editing. All authors read and approved the final version of the manuscript.

## Corresponding authors

Correspondence to Yingfei Ma.

## Competing interests

The authors declare no competing interests.

## Notes

### Competing Interest Statement

The authors have declared no competing interest.

