## Supplementary material for "Phage administration with repeated intravenous doses leads to faster phage clearance in mammalian hosts": sup: Supplementary material 0201bio archive.pdf

### Transmission Electron Microscopy Studies of Phages

Transmission Electron Microscopy studies were performed on the phage preparations and revealed a homogeneous population of phage PA_LZ7 and AB_SZ6 particles belonging to the *Myoviridae* and *Podoviridae* morphological groups, respectively. The phage PA39 reveals an icosahedral head structure of 73.6 nm and a contractile tail of 138.6 nm; the phage AB6 reveals an icosahedral head structure of 54.2 nm (see **Figure S1A**). The phage SE_SZW1 particles belonging to the *Siphoviridae* morphological group with an icosahedral head structure of 70.2 nm and a contractile tail of 158.6 nm[1].

### Genome Sequence Analysis of Phages

The sequence of phage AB_SZ6 has been deposited in the Genbank databases under accession number ON513429. The length of the phage's genome was 40,565-bp. Sequence-analysis indicated that it is affiliated to the *Podoviridae* family, *Caudovirales* order, *Friunavirus* genus. The sequence of phage PA_LZ7 has been deposited in the Genbank databases under accession number ON759747. The length of the phage's genome was 66,155-bp. Sequence-analysis indicated that it is affiliated to the *Myoviridae* family, *Caudovirales* order, *Pbunavirus* genus. The length of the phage SE_SZW1 genome was 45,881-bp. Sequence-analysis indicated that it is affiliated to the *Siphoviridae* family, *Caudovirales* order[1].

The linear genetic map of SE_SZW1, AB_SZ6 and PA_LZ7 can be seen in **Figure S1B**. Analysis of the phage whole genome shows that they possess a series of genes encoding common phage-related features, including DNA polymerase, DNA helicase, tail and head structure proteins. They also possess two genes encoding the host lysis protein, endolysin and holin. Importantly, the *in-silico* analysis did not reveal any putative virulence or antibiotic resistance or integrase sequences in the genome of these phages.

### Comparation of Plaque Assay and qPCR Analysis

The quantification for phage samples with a dilution series from $10^{10}$ to $10^2$ PFU/ml using plaque assay and qPCR analysis were evaluated. The result suggested the accuracy of qPCR analysis for phages in the range from $10^2$ to $10^{10}$ PFU/ml (**Figure S2A**).

### Phage Distribution in Organs with qPCR Analysis with A Single Dose in Rats

Phages accumulated largely in the spleen (7.83 ± 0.26 $Log_{10}$Copy/g) and the liver (6.97 ± 0.49 $Log_{10}$Copy/g) at 1h post-administration (**Figure S2B**). Similar to the active phage titer, a global decline in phage genome titer across all organs was observed within 72 h. Interestingly, the phage genome titer was ~2 $Log_{10}$Copy/g higher ($P$<0.001) than the active phage titer in the liver and lung at 72h post-administration (**Figure S2C**). This is likely due to the presence of the remaining DNA of the inactive phages.

#### PK of Phage SE_SZW1 with A Single Dose in Monkeys

At 5 min post-administration, the phage SE_SZW1 titer was 6.66 ± 0.51 and 7.02 ± 1.36 $Log_{10}$PFU/mL in the RLD and LD groups, respectively. At 24 h after IV administration, the active phage titer dropped to LLQ in both groups.

#### Tolerance Study of Phages with Repeated Doses in Healthy Animals

##### Clinical Sign and Blood Analysis

During the experimental period, no deaths or clinical signs related to the administration of phage SE_SZW1, AB_SZ6, and PA_LZ7 were observed. In addition, there were no significant differences in body mass between the phage-treated and the control group (**Table S4, S5, S6, and S7**). Furthermore, treatment with phage SE_SZW1 had no effects on the temperature, blood pressure, or electrocardiography in monkeys (**Table S7**).

Increased ($P$<0.0001) RWS was noted after the completion of injections (D7) of all three phages in the HD group in rats, however, this change was no longer observed following 14 days of recovery (D21) (**Figure S6**). There were no significant ($P$>0.05) differences in the relative weight of other organs in all tested animals. These phages had no significant hematologic, serum biochemical, immunological, or coagulation effects on the physiological parameters (**Table S4, S5, S6, and S7**).

##### Histology Analysis

Slightly extramedullary hematopoiesis (EMH) was observed in the spleen samples from the phage SE_SZW1 HD group (3 out of 5 rats), phage PA_LZ7 HD group (2 out of 6 rats), and LD group (2 out of 6 rats) after 7 administrations (D7) (**Figure S7**). No more EMH was detected following the completion of the experiment (D21). No clinically significant histopathologic findings were noted in any of the brain, heart, lung, kidney, liver, thymus, bronchus, or gastrointestinal tract samples (**Figure S8**). No clinically significant histopathologic findings were detected in any samples from rat models with phage AB_SZ6 and monkey models with phage SE_SZW1.

### Methods

#### Bacterial Strains

*Salmonella typhimurium* strain SL7207 and *P. aeruginosa* strain PAO1 were from laboratory collections. *A. baumannii* clinical isolate was obtained from routine microbiological cultures of clinical samples. Bacterial strains were maintained in Luria-Bertani (LB) broth (Huankai Microbiol, Guangzhou, China) and stored in 15% glycerol at -80°C.

### Isolation of Phages

Phages were isolated from various environmental samples by using routine isolation techniques, as previously described[2]. Briefly, bacterial strains were used to isolate and propagate pathogen-specific phages. Following isolation, the phages were triple plaque-purified on their respective host bacterium. Finally, small-scale phage amplification on their corresponding host bacterium was performed to prepare the phage library, which was subsequently stored at 4°C until required.

### Propagation and Purification of Phages

Phages were propagated on their corresponding host bacterium using 1 liter LB broth, yielding lysates with titers of around $10^{10}$ PFU/mL. Lysates were filtered using 0.22-μm filters and then concentrated with the cross-flow filtration method by using a 100 kDa membrane (Jiuling, Hangzhou, China). Prepare a cesium chloride step density gradient of 1.7, 1.5, and 1.3 g $cm^{-3}$ with SM buffer (100 mM NaCl, 8 mM MgSO4, 50 mM Tris-HCl, pH 7.5) in an open-top ultraclear round-bottom tube, layer the phage concentrate on the top and ultracentrifuge at 24,000 rpm/min for 2 h at 4°C. The visible phage band was collected (approximately 1.5 mL), then dialyzed using a Spectra/Por RC 6 Dialysis membrane (MWCO 25 K Da, Sangon, Shanghai, China) in SM buffer to remove cesium chloride. Phages were then sterilized through 0.22-μm filters. Then phages were titrated and evaluated for endotoxin with an End-point Chromogenic Endotoxin Test Kit (Bioendo, Xiamen, China). Phage preparations were subsequently stored at 4°C until required.

### Phage DNA Extraction, Genome Sequencing, and Assembly

Phage particles were precipitated with 10% polyethylene glycol 8,000 (PEG 8000) at 4°C overnight, centrifuged at 10,000 g for 15 min, and subsequently suspended in SM buffer. Then the concentrated phage particles were treated using DNase I and RNase A (New England BioLab, Massachusetts, USA) to remove bacterial nucleic acids, genomic phage DNA was extracted with MiniBEST Viral RNA/DNA Extraction Kit (Takara, Beijing, China) following manufacturer's protocol. Whole genome sequencing was performed at the Tianjing Sequencing Center (Novogene, Beijing, China) using the Illumina HiSeq system (Illumina Inc., San Diego, CA, USA). Reads were assembled with SOAPdenovo2[3]. The resulting contigs were uploaded into the RAST server using the RASTtk annotation workflow[4]. Putative functions of the ORFs were further identified with Blast-P based on amino acid sequences. A linear map of the phage was depicted using the Proksee Server (https://proksee.ca/).

### Transmission Electron Microscopy

20 μl of phage suspension with a pipet-gun were dropped onto the copper grid with carbon film for 3-5 min, and then used filter paper to absorb the excess liquid. 2% phosphotungstic acid on the copper grid was dropped to stain for 1-2 min, filter paper was used to absorb excess liquid and dried at room temperature. The cuprum grids were observed under Transmission Electron Microscopy (HT7800, Hitachi, Tokyo, Japan).

**Animals**

Male and female Sprague-Dawley rats (6-8 weeks old; males weigh around 250g; females weigh around 200g) were purchased from Pengyue Experimental Animal Breeding Co. (Jinan, Shangdong, China). Male and female cynomolgus monkeys (*Macaca fascicularis*; 5-7 years old) were purchased from Zhongkelingrui Biological Technology Co. (Beijing, China). See **Table S4, S5, S6, and S7** for more information, including the sex, weight, and age of the animals. The animals were housed under a 12-h dark-light cycle with free access to a standard diet and water. All animal experiments were approved by the Shandong Academy of Pharmaceutical Sciences Animal Ethics Committee.

**Phage Administration**

The phage preparation was diluted with PBS and administered to rats through the tail vein. The phage preparation was diluted with 0.9% saline and administered to cynomolgus monkeys through the superficial vein of the left hind limb.

**PK and Biodistribution Study**

Healthy female rats received phage SE_SZW1, PA_LZ7, and AB_SZ6 once daily by intravenous (IV) injection at $5*10^9$ or $5*10^{10}$ PFU/kg per day in a volume of 10 mL/kg for 7 days (n=5 per dose group). Control groups were treated with sterile PBS only (n=5). The jugular vein of rats was cannulated for blood collection. Blood samples were collected at 5 and 30 min, and 1, 2, 4, 6, and 24 h following the first intravenous administration. For animals that received phages SE_SZW11 and AB_SZ6, the blood samples were collected at 5 min following 3, 5, and 7 IV doses as well. The single-dose biodistribution study was performed in healthy female rats following an IV bolus of phage at dosages of $5*10^9$ or $5*10^{10}$ PFU/kg (n=15 per dose group). At 1, 24, or 72 h, rats were humanely killed (n=5 per time point) and tissues (i.e., lung, kidney, spleen, brain, and liver) were harvested and homogenized in SM buffer. The biodistribution study of repeated doses was performed in healthy female rats following 3 IV bolus of phage SE_SZW1 at dosages of $5*10^{10}$ PFU/kg (n=15). At 1, 24, or 72 h post the 3rd dose, rats were humanely killed (n=5 per time point) and spleen and liver were harvested and homogenized in SM buffer.

20 healthy female rats were divided into 4 groups (n=5) to check whether phage could induce enhanced

nonspecific phage clearance. Animals received 2 doses of phages SE_SZW1, AB_SZ6, and PA_LZ7 at a dosage of $5*10^{10}$ PFU/kg or PBS and consecutively 1 dose of phage SE_SZW1 in 3 days. Blood samples were collected at 5 and 30 min and 1 h following the last intravenous administration.

Healthy monkeys received phage SE_SZW1 once daily by IV administration at dosages of $10^9$ or $5*10^9$ PFU/kg per day in a volume of 1 mL/kg for 14 days (2 male and 2 female monkeys for $10^9$ PFU/kg group; 3 male and 3 female monkeys for $5*10^9$ PFU/kg group). The control group was treated with sterile 0.9% saline only (1 male and 1 female monkey). Blood samples were collected at 5 and 30 min, and 1, 2, 4, 6, and 24 h following the first intravenous administration. Blood samples were collected at 5 min following 3, 7, 11, and 14 IV doses as well. The distribution study in healthy monkeys was performed following 14 IV doses of phage SE_SZW1 at $10^9$ and $5*10^9$ PFU/kg (2 male and 2 female monkeys per dose group). At 24 h, monkeys were humanely killed and tissues (i.e., lung, kidney, spleen, brain, and liver) were harvested and homogenized in SM buffer.

For the PK study, regression analysis on the phage titer in plasma over time was performed using non-compartmental analysis. Semi-log plots were constructed of the individual plasma phage concentration versus time profiles for each subject. Area under the curve (AUC) values were calculated by log-linear trapezoidal integration using the Phoenix WinNonlin 8.1 (Certara, Princeton, NJ, USA). $\lambda z$ was the calculated slope of the terminal portion of the log plasma concentration versus the time curve. Extrapolation of the AUC from the last measured plasma concentration to infinity was calculated as Cpn/ $\lambda z$ where Cpn is the last measured plasma concentration. Plasma clearance (CL) was calculated as Dose/$AUC_{inf}$. At least three-time points in the terminal phase were required to calculate $\lambda z$.

**Phage Quantification in Animal Samples**

Samples were generally analyzed for phage enumeration with the double agar overlay method as follows: 20 µl plasma was transferred to a 1.5 mL Eppendorf tube and 0.48 mL SM buffer was added, after vortexing, samples were passed through 0.22 µm filters; tissues were harvested and homogenized in SM buffer with ration 1:5 (m/v), then samples were passed through 0.22 µm filters; following 10-fold serially diluted, 10 µl added to their corresponding host bacterium agar overlays. The lower limit of phage quantification (LLQ) was 5,000 and 1,200 PFU/mL (i.e., 1 PFU per plate) for plasma and tissue samples, respectively. Samples were also analyzed with the quantitative PCR (qPCR) method for phage SE_SZW1[5]. The yield and purity of isolated DNA were measured by a Qubit 4 Fluorometer (ThermoFisher Scientific, Merelbeke, Belgium). The primers sequence targets the gene encodes major coat protein of phage SE_SZW11 (primers: forward, 5′-ACCAAAGACTTGGTAGCCAACTC-3′ and reverse, 5′- TCGCGCCAGTAATCTTTCGTT-3′). qPCR was performed with AceQ qPCR SYBR Green Master Mix (Vazyme, Nanjing, China) using the qTower$^3$ system

(Analytik jena, Jena, Germany).

### Endotoxin Study

Endotoxin used in this manuscript is extracted from *Salmonella typhimurium* SL7207 by hot aqueous-phenol extraction method[6]. Endotoxin was diluted with PBS to the same level of the SE_SZW1 HD group and administered to rats through the tail vein.

10 healthy female rats were divided into 2 groups (n=5) to check whether endotoxin could induce increased pro-inflammatory effect or enhanced non-specific phage clearance. Animals received 2 doses of endotoxin or PBS and consecutively 1 dose of phage SE_SZW1 in 3 days. Blood samples were collected at 1 and 24 h for cytokines measurement following the first intravenous administration. Blood samples were collected at 5, 30 min and 1 h following the last intravenous administration for phage quantification.

### Cytokines Quantification

The cytokines level of rats was evaluated by V-PLEX Proinflammatory Panel 2 Rat Kit with measurements by Meso QuickPlex SQ120 following the manufacturer's instructions (Meso Scale Discovery, Rockville, MD, USA). TNF-α, IL-6, IL-10, IFN-γ, IL-2, IL-1β, IL-8, MIP-1a, KC/GRO, and GSF level of monkeys was measured by Elisa Kit following the manufacturer's instructions (Enzyme-linked Biotechnology Co., Shanghai, China).

### Adaptive Immune Response Study

Following phage administration, animals underwent a 14-day recovery period (without phage administration). Rats' blood samples were collected on D1 (prior phage injection), 3, 5, 7, 10, 12, 15, and 21 for the immune response study; monkeys' blood samples were collected on D1 (prior phage injection), 3, 7, 11,14, 21 and 28 for immune response study.

### Phage Neutralization Assays

Phage neutralization assay was performed as previously described with several modifications[7]. Plasma samples collected from phage SE_SZW11 and AB6 treated groups were incubated with corresponding phages. 1 μl of plasma was incubated with 9 μl of SM buffer (containing about $4.5*10^5$ PFU phage) at room temperature for 24 h. After incubating the plasma-phage mixtures, ten-fold serial dilutions were made and 5 μl of each dilution was spotted onto agar containing corresponding host bacteria. Plates were incubated at 37°C overnight. Each test was investigated in triplication. The controls are the plasma collected on day 1 before phage injection.

**Western Bolt**

Western blot analysis was performed as previously described with several modifications[7]. Phage protein samples were prepared by mixing highly purified phages ($4*10^{10}$ PFU per well) with 5×SDS-PAGE loading buffer and boiling at 100°C for 10 minutes. Phage proteins were separated by electrophoresis on 4-20% SDS-PAGE gradient gel at 140V for 55 minutes and transferred to the PVDF membrane at 15V for 20 minutes. The membrane was blocked in TBST (TBS with 0.1% Tween 20) +5% milk for 5 h at 37°C and then washed 3 times with TBST. The membrane was incubated overnight with heat-inactivated plasma diluted 1:1000 in TBST+0.5% milk at 4°C. Plasma was removed, and the membrane was washed 3 times with TBST and then, incubated for 2 h with the secondary antibody diluted 1:12500 in TBST+0.5% milk at 37°C. Secondary antibodies included: HRP conjugated goat anti-rat IgG Fc preadsorbed (catalog no. ab97090; Abcam, Cambridge, UK) and HRP conjugated goat anti-monkey IgG gamma (catalog no. FNSA-0123; FineTest, Wuhan, China). Finally, the secondary antibody was decanted and the membrane was washed 3 times in TBST. The membrane was incubated for 1min with super ECL detection reagent in the dark and imaged with UVP ChemStudio (Analytik jena, Jena, Germany).

**Protein Identification by Mass Spectrometry**

After SDS-gel electrophoresis, part of the gel was excised and Coomassie-staining was used to obtain visible protein bands. Protein bands that were identified by the western bolt were excised and in-gel digested with trypsin. The resulting peptides were analyzed by liquid chromatography-tandem mass spectrometry and searched using Maxquant software (version 2.0.1). The protein entry with the highest intensity and corresponding molecular weight was regarded as the target protein.

**Phage Tolerance Study**

The tolerance study was performed using the previously described method with several modifications[8]. Three groups of 36 rats (18 males and 18 females) were used for phage PA_LZ7. Six healthy animals of each gender received phage PA_LZ7 once daily by IV injection at dosages of $5*10^9$ or $5*10^{10}$ PFU/kg per day in a volume of 10 mL/kg for 7 days. Control groups were treated with sterile PBS only. 3 animals of each gender from each group were sacrificed following 7 IV phage injections, the rest animals underwent a 14-day recovery period (without phage administration).

Three groups of 30 healthy female rats were used for phage SE_SZW1 or AB_SZ6. 5 healthy animals received phage once daily by IV injection at dosages of $5*10^9$ or $5*10^{10}$ PFU/kg per day in a volume of 10 mL/kg for 7 days. Control groups were treated with sterile PBS only. 5 healthy animals from each group were sacrificed following 7 IV phage injections, the rest animals underwent a 14-day recovery period (without phage administration).

Monkeys used for the PK study were used for the safety study as well, 2 animals of each gender from phage administration groups were sacrificed following 14 IV phage injections, and the rest animals underwent a 14-day recovery period (without phage administration).

During the whole period of the experiment, the animals were observed daily to detect any signs of toxicity.

**Clinical Signs**

The animals were observed continuously for any clinical signs or mortality for the first 6 hours after the first administration, next they were examined once daily following administration, then they were checked once per week during the recovery period. The animals were observed for clinical signs, including changes in appearance, posture, movement, urine, body surface, and fluid secretion/excretion, throughout the experimental period.

The monkeys were additionally examined for temperature, blood pressure (BP), and electrocardiogram (ECG). Animals' rectal temperature was measured twice before administration (D-5 and D-1), next they were examined at 2 and 24 h following administration (D1, 4, 7, and 14). Then they were checked once per week during the recovery period. Animals' BP and ECG were measured once before administration, next, they were examined once following administration (D13), and then they were checked once during the recovery period (D26). The monkeys were anesthetized using IV injections with Zoletil (Virbac, Carros, France), then ECG and BP assessments proceeded with a PACK4G Jacket telemetry (emka Tech, Sterling, VA, USA).

**Body Mass**

Animals were weighed before administration (D-1) and once weekly thereafter.

**Food Consumption**

Food consumption was measured on the initial day of treatment, and then once weekly during the experimental period for rats. The amounts of food placed in each cage were measured, as were the remaining amounts the next day, to calculate the daily food consumption (g/rat/day). Food consumption was measured daily for monkeys based on the remaining food for each animal, the animal's food intake was estimated as 100%, more than 75%, 75%, 50%, 25%, and 0%.

**Histopathologic Analysis**

Before necropsy, all surviving animals were fasted overnight (for 16–20 h), anesthetized using intraperitoneal injections with Zoletil (Virbac, Carros, France) / xylazine for rats; and using IV injections with Zoletil / propofol for monkey. After anesthesia was confirmed, exsanguination was performed to euthanize the animals. An initial inspection was then made of the body surface, subcutis, head, and all internal organs of the abdominal and thoracic cavities. Next, the brain, liver, lung, heart, spleen, kidneys, thymus, prostate gland, testes, epididymis, ovaries, and uterus were removed and examined separately. Then the brain, liver, lung, heart, spleen, kidneys, and thymus were weighed.

Histological examination of organs was done as described previously[9]. Briefly, tissues were fixed with 10% formalin and embedded in paraffin wax. Serial sections of 3 μm thickness were cut using a microtome, de-paraffinized, rehydrated and stained with Hematoxylin and Eosin (HE). For histological changes, the tissue sections were examined under the Nikon E100 microscope (Nikon, Tokyo, Japan).

**Hematology and Serum Biochemistry**

Blood samples were collected from the abdominal aorta of all rats and the peripheral vein of all monkeys scheduled for necropsy under deep anesthesia. Before the blood was collected into tubes, containing potassium EDTA as an anticoagulant, for hematologic analysis, using a hematology analyzer (XT-2000 iV, Sysmex Corporation, Kobe, Hyogo, Japan). About 1.8 mL of blood was collected from each animal into tubes containing sodium citrate and was analyzed using an automated CA1500 blood coagulation analyzer (Sysmex Corporation, Kobe, Hyogo, Japan). Approximately 2 mL of blood was collected from each animal into tubes for biochemical analysis, which employed an automated 7180 clinical chemistry analyzer (Hitachi, Tokyo, Japan).

**Total Immunoglobulin Measurements**

IgG, IgM, IgA, C3, and C4 level of rats was measured by Elisa Kit following the manufacturer's instructions (Enzyme-linked Biotechnology Co., Shanghai, China).

**Flow Cytometry**

Whole blood samples were incubated with antibody cocktails (1:100), for 20 min at 4°C. Antibody combinations: FITC-conjugated anti-rat CD45RA, PE-conjugated anti-rat CD161, APC-conjugated anti-rat CD3; FITC-conjugated anti-rat CD4, PE-conjugated anti-rat CD8a, APC-conjugated anti-rat CD3 (BioLegend, San Diego, CA, USA). Added lyse buffer to mixed solution, after vortexing, incubated for 15 to 60 minutes at room temperature, protected from light. Samples were washed 3 times with PBS, resuspended in PBS, and kept at 4°C up to analysis. Flow cytometry was performed on Guava easyCyte 6-2L HT Flow Cytometer (Luminex Corporate, Austin, TX, USA), and data were analyzed with Guavasoft (3.1.1) software (Luminex Corporate, Austin, TX, USA).

### Data analysis

Comparisons were performed by one-way ANOVA with Bonferroni's multiple comparisons test and Student's t-test except for special explanation. All statistical analyses were performed using Prism 7.04 (GraphPad, San Diego, CA, USA), and differences with $P< 0.05$ were considered statistically significant.

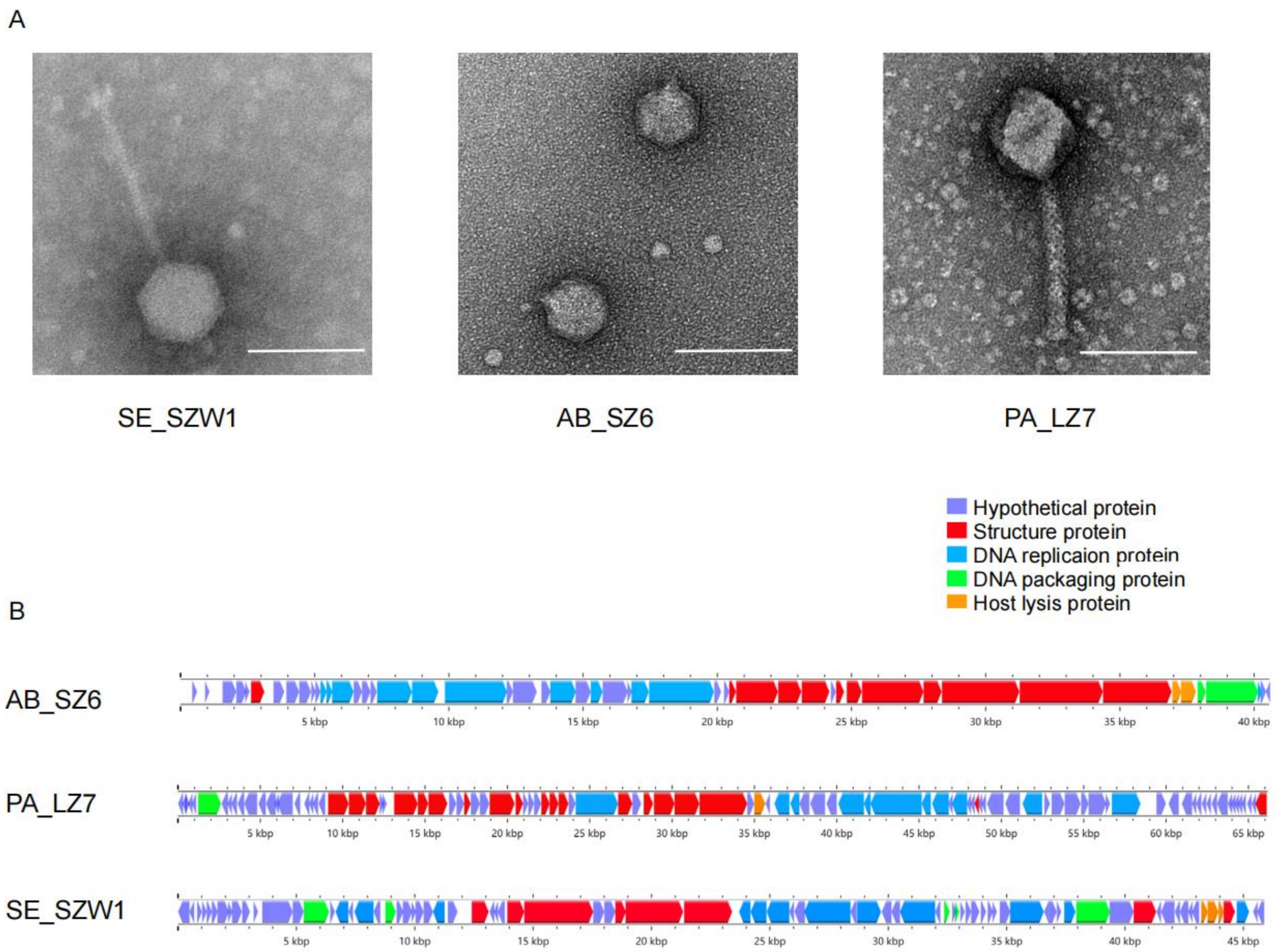


**Figure S1. Characterization of the phage used in this study.** (A) Transmission electron micrographs of the phage. The scale bar represents 100 nm. (B) Genome map of the phage. The color of the ORFs refers to five modules: phage structure, red; host lysis, orange; DNA packaging, green; DNA replication, light blue; and hypothetical proteins, purple.

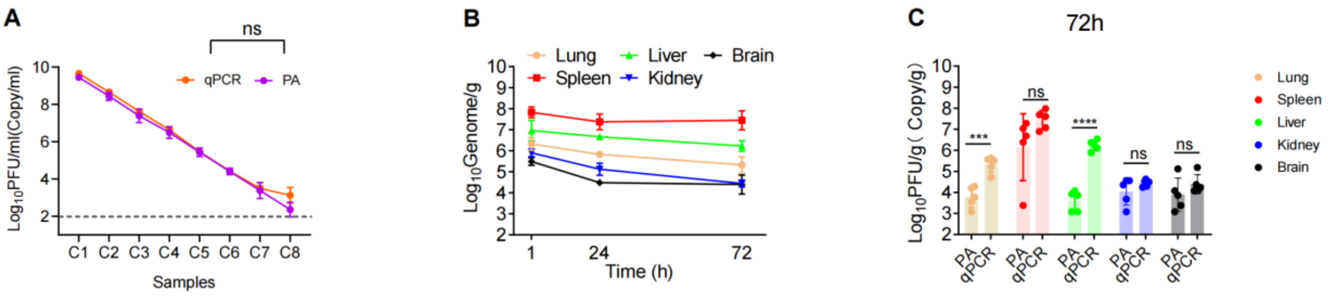


**Figure S2. Phage enumeration with qPCR analysis.** (A) Comparison of phage quantification for dilution series of phage SE_SZW1 stock using plaque assay and qPCR method. The X-axis represents the dilution of the phage stock, and Y-axis represents the concentrations of active phage (PFU/ml) or phage genome (copy/ml) obtained with plaque assay or qPCR method, respectively. The limit of detection is represented by the grey dashed line. Mann-Whitney test was performed in this assay (n=3; ns: non-significant). (B) Biodistribution of phage SE_SZW1 genome titer following single IV administration in the HD group. Phage genome titer is expressed as copy per g. Phage genome titer was determined by the qPCR method, each symbol representing the means with sd (n=5). (C) Comparations of active phage titer by plaque assay and phage genome titer by the qPCR method for phage SE_SZW1 in the HD group at 72h post-administration. PA: plaque assay; qPCR: qPCR analysis. Data are presented as means with sd (n=5; *, $P<0.05$; **, $P<0.01$; *** $P<0.001$; **** $P<0.0001$; ns: no significance).

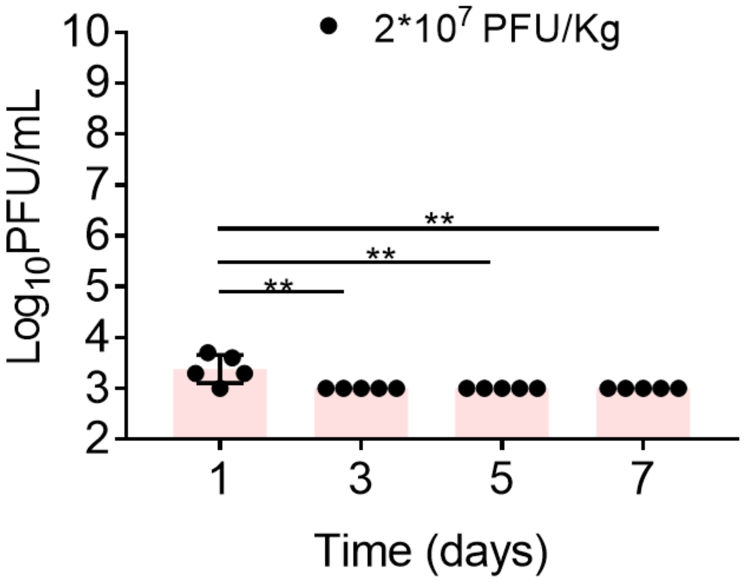


**Figure S3. Active phage titer at 5 min in plasma after different IV injections with phage SE_SZW1 at 2*$10^7$ PFU/kg.** Phage titer was obtained by plaque assay, each symbol represents the means with sd (n=5). The LLOQ is 1000 PFU/mL (**, *P*<0.01).

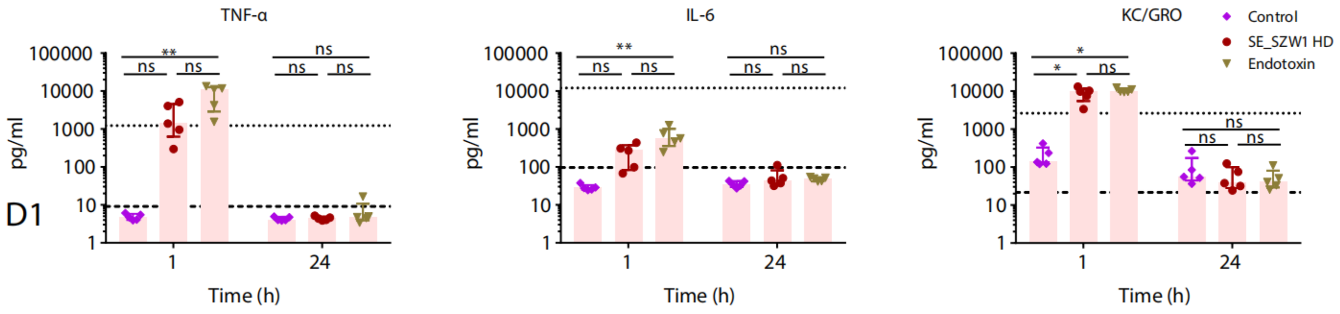


**Figure S4. Cytokine profile in plasma of rats following endotoxin administration.** Cytokine concentrations in plasma were measured at 1 and 24h after administration with *Salmonella* endotoxin or PBS. Endotoxin was diluted with PBS to the same level of the SE_SZW1 HD group. Data are presented as median with IQR. Comparisons were performed exclusively within the same time point (n=5; ** $P<0.01$; ns: no significance). Analysis was performed by Kruskal-Wallis with Dunn’s test.

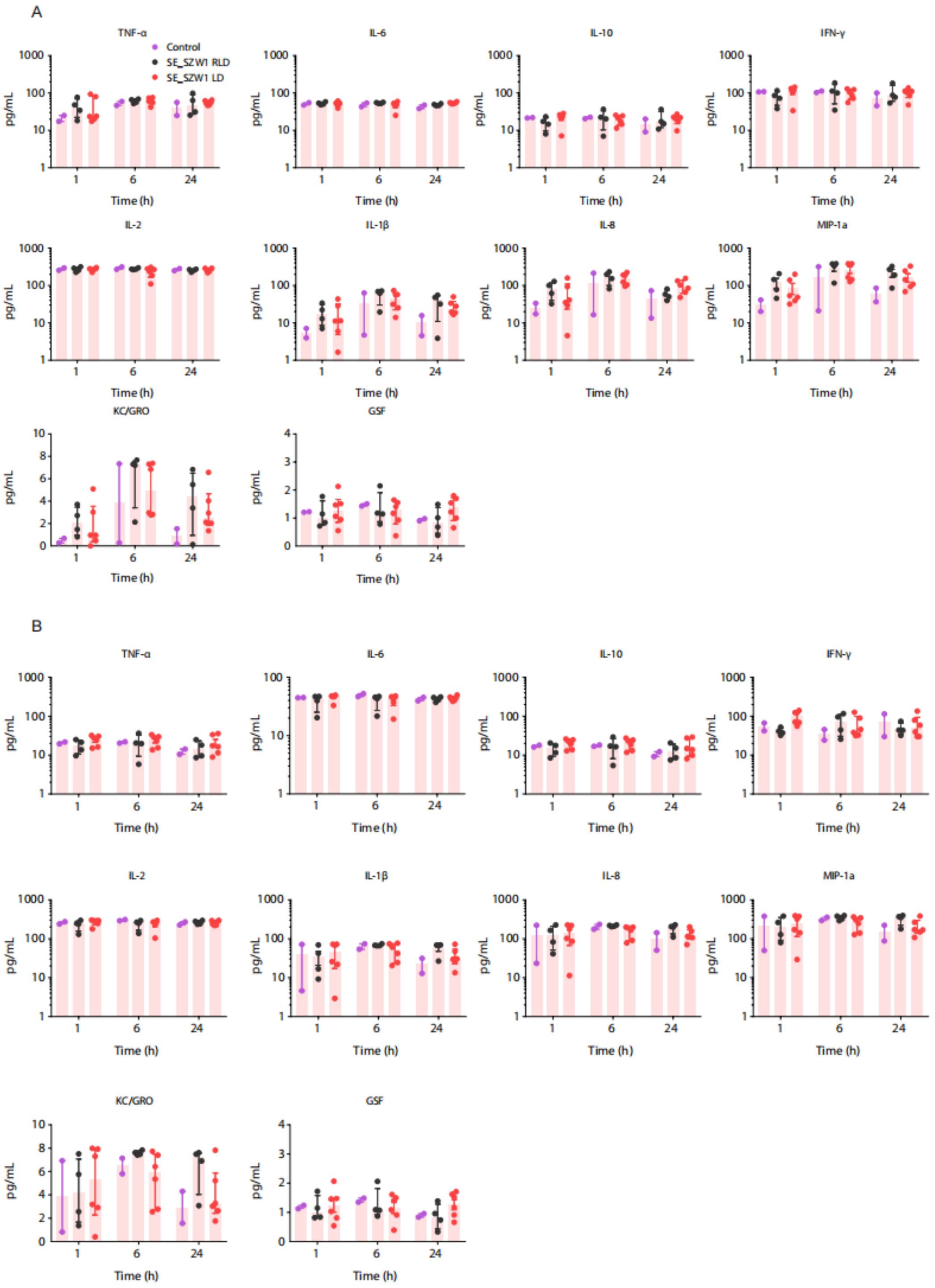


**Figure S5. Cytokine profile in plasma of monkeys following phage administration.** Cytokine concentrations in plasma were measured at 1, 6 and 24 h with phage SE_SZW1 following a single dose or 14 repeated doses on D1 (A) and D14 (B). Monkeys received 0.9% saline (Control, n=2) or were treated by phages with $10^9$ PFU/kg (RLD, n=4) or with $5*10^9$ PFU/kg (LD, n=6). Data are presented as median with IQR. Comparisons were performed exclusively within the same time point (ns: no significance).

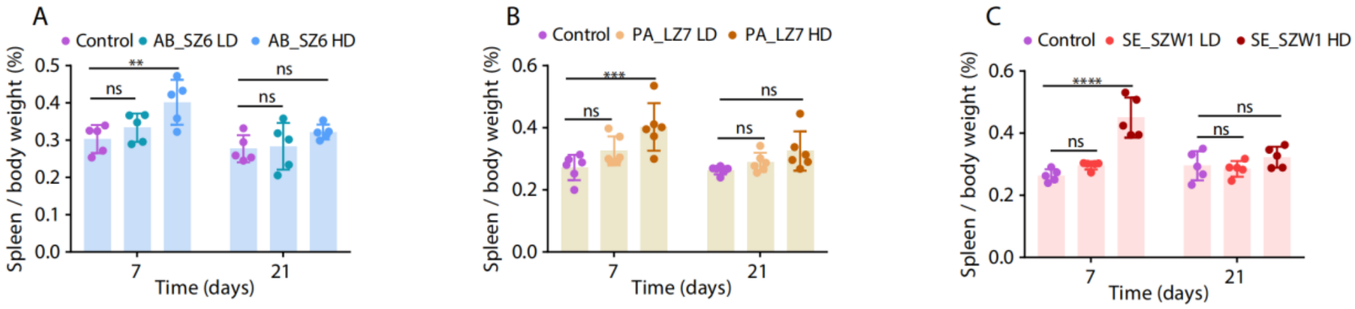


**Figure S6. The relative weight of the spleen changes in rats.** The relative weight of the spleen changes in rats following 7 doses and a 14-day recovery period in both LD and HD groups with phage AB_SZ6 (A) and PA_LZ7 (B), and (C). Data are presented as means with sd (n=5; ** *P*<0.01; *** *P*<0.001; **** *P*<0.0001; ns: no significance).

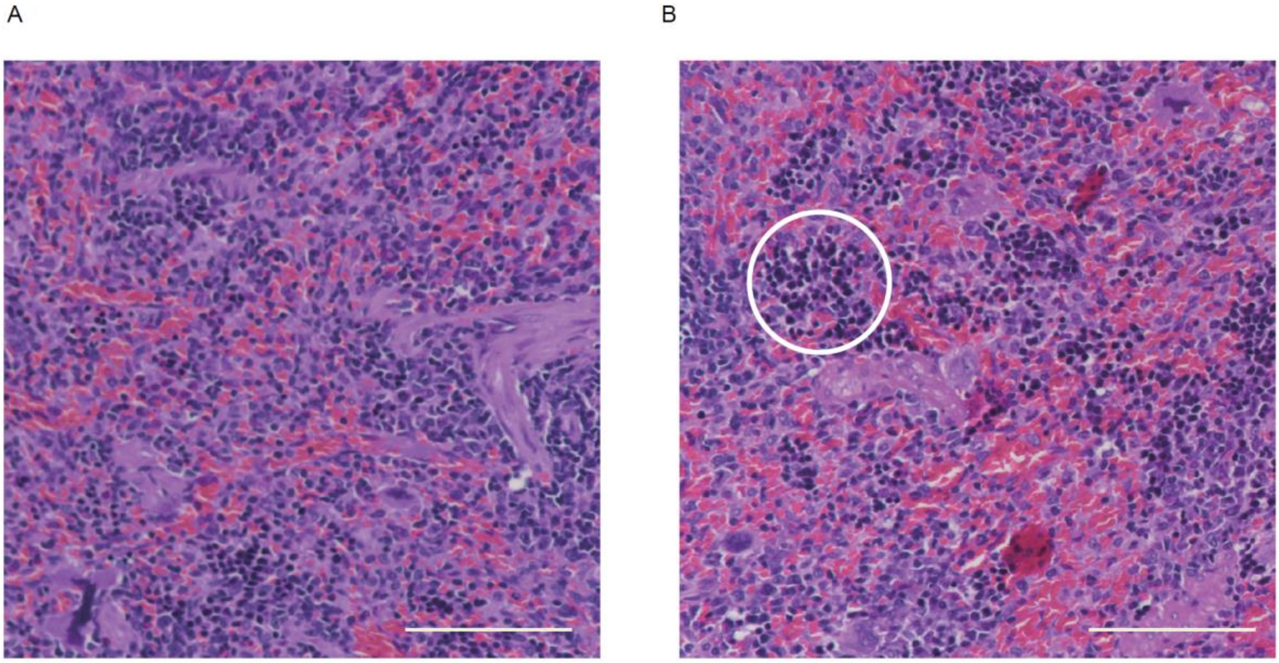


**Figure S7. Representative hematoxylin and eosin (HE) stained section of the spleen:** (A) normal structure; (B) many erythroid precursors (white circle) appear in the red pulp. The scale bar represents 100 $\mu$m.

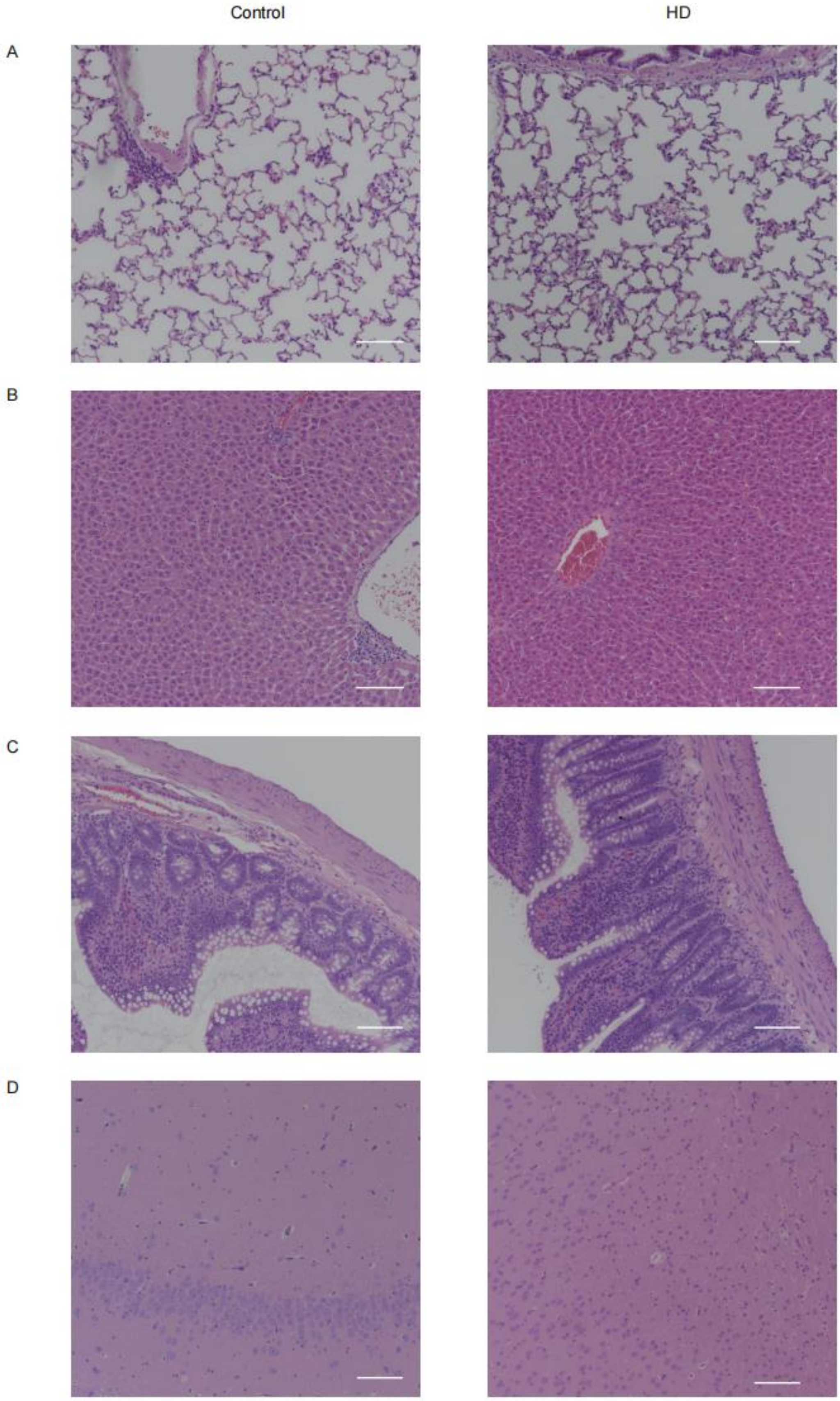


1

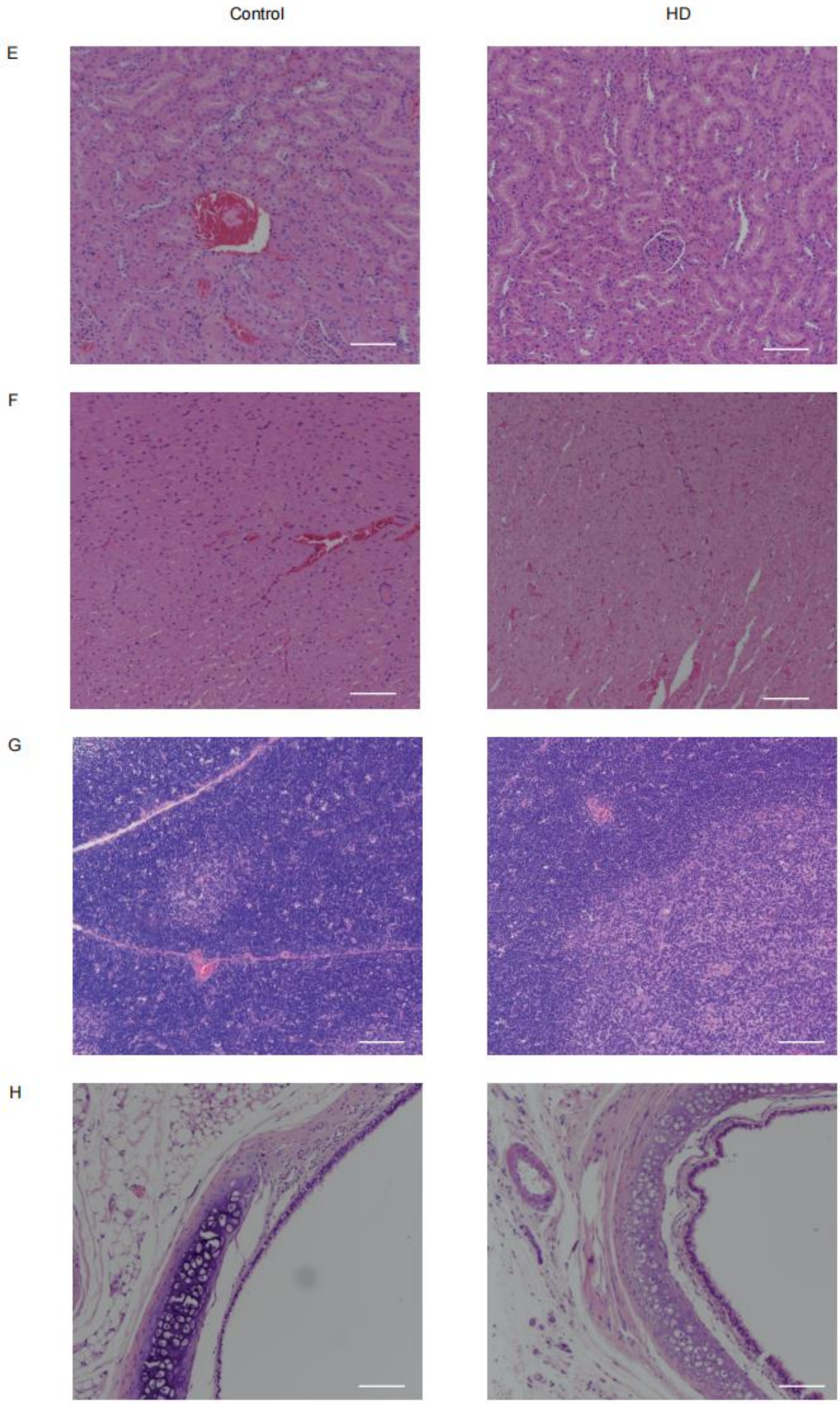


**Figure S8. Representative HE stained section of the normal structure of other organs in animals with PBS or phage PA_LZ7 with HD:** (A) lung, (B) liver, (C) ileum, (D) brain, (E) kidney, (F) heart, (G) thymus and (H) bronchus. The scale bar represents 100 $\mu$m.

| Phage | Phage family | Bacterial host | Phage titer (PFU/mL) | Endotoxin concentration (EU/mL) | Endotoxin unit per $10^9$ PFU |
|---|---|---|---|---|---|
| SE_SZW1 | *Siphoviridae* | *Salmonella typhimurium* strain SL7207 | $4*10^{12}$ | $1.5*10^4$ | 3.75 |
| AB_SZ6 | *Podoviridae* | *A. baumannii* strain AB-SZ01 | $10^{12}$ | $2*10^3$ | 2 |
| PA_LZ7 | *Myoviridae* | *P. aeruginosa* strain PAO1 | $2*10^{11}$ | $4*10^3$ | 20 |

**Table S1. Information of phages used in this study.** For each phage, taxonomical family and bacterial host strain are presented, as well as the titer of the phage and level of endotoxin.

| Parameter | SE_SZW1 HD | PA_LZ7 HD | AB_SZ6 HD | SE_SZW1 LD | PA_LZ7 LD | AB_SZ6 LD |
|---|---|---|---|---|---|---|
| $C_{max}$ $Log_{10}$(PFU/mL) | 8.72±0.39 | 8.62±0.41[b] | 5.47±0.21[c] | 7.60±0.29[d] | 6.18±0.97[e] | 4.61±0.92[f] |
| CL (L/h/kg) | 0.09±0.04 | 0.33±0.21[b] | 280.47±78.38[c] | 0.16±0.16[d] | 6.16±3.74[e] | 56.56±12.36[f] |
| $AUC_{0\text{-}n}$ $Log_{10}$(h*PFU/mL) | 8.81±0.22[a] | 8.23±0.24[b] | 5.27±0.13[c] | 7.63±0.33[d] | 6.11±0.62[e] | 4.95±0.09[f] |

**Table S2. PK parameters of three phages in rats.** $C_{max}$, maximum observed plasma concentration; CL, plasma clearance; $AUC_{0\text{–}n}$, area under the concentration-time curve from time 0 to infinity. Data are presented as mean with sd ([a]: SE_SZW1 HD *vs* PA_LZ7 HD, $P$<0.05; [b]: PA_LZ7 HD *vs* AB_SZ6 HD, $P$<0.05; [c]: SE_SZW1 HD *vs* AB_SZ6 HD, $P$<0.05; [d]: SE_SZW1 LD *vs* PA_LZ7 LD, $P$<0.05; [e]: PA_LZ7 LD *vs* AB_SZ6 LD, $P$<0.05; [f]: SE_SZW1 LD *vs* AB_SZ6 LD, $P$<0.05; $P$<0.05).

| Parameter | LD | RLD |
|---|---|---|
| $C_{max}$ $Log_{10}$(PFU/mL) | 5.73±0.89[a] | 5.99±0.57 |
| CL (L/h/kg) | 2.11±3.65 | 0.60±0.62 |
| $AUC_{0\text{-}n}$ $Log_{10}$(h*PFU/mL) | 6.87±0.72 | 6.37±0.41 |

**Table S3. PK parameters of phage SE_SZW1 in monkeys.** $C_{max}$, maximum observed plasma concentration; CL, plasma clearance; $AUC_{0\text{–}n}$, area under the concentration-time curve from time 0 to infinity. Data are presented as mean with sd ([a]: SE_SZW1 LD in rats *vs* SE_SZW1 LD in monkeys, $P$<0.05).
